# MRI-Based Blood Clot Phenotyping: An In Vitro Study

**DOI:** 10.64898/2026.04.14.718500

**Authors:** Grace N. Bechtel, Ayesha B. Das, Juliette Noyer, Adam M. Bush, David A. Hormuth, Thomas E. Yankeelov, Edward Castillo, Steven Warach, Jan Fuhg, Jonathan I. Tamir, Hamidreza Saber, Manuel K. Rausch

## Abstract

**Background and Purpose:** Neurointerventional outcomes depend on clot composition and may be influenced by clot contraction. Thus, a priori identification of clot composition and contraction could inform procedural strategies and improve outcomes. The goal of our work is to conduct an in vitro test to determine whether MRI can reliably predict both clot composition and contractile state.

**Materials and Methods:** To this end, we prepared blood clots spanning clinically observed compositions (0-80% red blood cells (RBCs)) in both contracted and uncontracted states. Contraction was controlled by coagulating blood with or without thrombin. We imaged these clots using quantitative, clinical, and investigational MRI sequences. Using these data, we then determined whether MRI signal intensities, quantitative parameters, and radiomic features capturing intensity and texture patterns can (i) predict clot hematocrit and (ii) classify clots by composition (RBC-rich vs. fibrin-rich) and contraction state.

**Results:** Quantitative MRI parameters (T1, T2, ADC) decreased with increasing hematocrit (R^2^ = 0.56–0.85, p < 0.001), while signal intensities from clinical sequences showed weaker correlations (R^2^ = 0.46–0.62, p < 0.001). Radiomic models predicted hematocrit with performance comparable to MRI parameters. When applied to classification, radiomic features accurately discriminated RBC-versus fibrin-rich clots, with AUCs exceeding 0.90 across nearly all sequences. In contrast, classification of contraction state showed greater variability in AUCs across sequences but remained high for quantitative T1 and T2 values (AUCs up to 0.88). Trends were consistent across clots coagulated with and without thrombin. Pooling features across sequences did not outperform the best individual sequence for either regression or classification.

**Conclusions:** We demonstrate that MRI-based radiomic analysis quantitatively characterizes clot composition and contraction in vitro. These findings support the feasibility of using MRI for pre-interventional clot phenotyping, with potential to inform thrombolytic and mechanical thrombectomy strategies. Thus, in vivo studies validating these results are warranted.

## INTRODUCTION

Acute ischemic stroke is a leading cause of death and disability worldwide^1,2^. Its treatment relies on rapid restoration of blood flow, which is accomplished via intravenous thrombolysis, mechanical thrombectomy, or both^3^. In all cases, treatment success depends on the composition of the occluding clot^4^. For instance, clots rich in red blood cells (RBCs) are generally softer and more porous, which leads to greater responsiveness to thrombolytic agents and easier aspiration^5^. In contrast, fibrin-rich clots are stiffer and denser, increasing resistance to lysis and reducing the likelihood of aspiration success^6^. Beyond composition, clots differ in their degree of contraction^7^. Contraction increases clot density and stiffness, which may also affect treatment success^8^. Thus, characterizing clot composition and contraction prior to intervention could meaningfully inform treatment strategy.

Despite its potential clinical importance, pre-interventional assessment of clot composition is currently limited. That is, conventional imaging primarily provides indirect markers^9^. For example, the hyperdense artery sign on CT and susceptibility-related blooming on MRI are used as markers of high RBC content^10^. However, both measures are qualitative, sequence-dependent, and inconsistently present^9^. Although emerging approaches such as dual-energy or spectral-detector CT show promise^11^, quantitative and reproducible imaging biomarkers for clot composition remain lacking.

MRI is increasingly used in acute stroke imaging^12^. This is partly due to its superior tissue characterization, particularly for detecting early ischemic changes^13,14^. However, even standard clinical stroke protocols were not designed to characterize the occluding clot^15^, and the sensitivity of different MRI sequences to clot composition and contraction remains incompletely understood. This includes investigational research sequences (e.g., ultra-short echo time) and quantitative mapping techniques (e.g., T1, T2, and ADC mapping)^16^. Although these approaches are not routinely used in acute stroke workflows due to longer scan times or limited clinical validation^17^, they may provide valuable information about clot composition and contraction. As a result, it remains unclear which clinical, quantitative, and investigational MRI approaches are most informative for clot phenotyping.

Radiomic analysis may provide a complementary approach for extracting information from medical images^18^. Specifically, radiomics quantifies subtle spatial and textural features beyond what is visually apparent^19^. In fact, radiomics has already shown promise for stroke imaging, particularly on CT. In these cases, radiomics has been used to predict clot composition and mechanical thrombectomy outcomes^20–23^. In contrast, although radiomics has been applied to MRI in other stroke contexts (e.g., prognosis prediction), its use specifically to characterize clot composition and contraction has not been tested.

In this study, we aim to determine the relationship between clot composition and MRI signal using a controlled in vitro model. We image both contracted and uncontracted clots, with RBC content ranging from 0 to 80%, using quantitative MRI sequences, standard clinical stroke sequences, and investigational sequences. We assess how MRI-based radiomic analysis can (i) predict clot hematocrit and (ii) classify clots by RBC-rich versus fibrin-rich composition and contraction state. Thus, this work ultimately seeks to demonstrate the feasibility of MRI-based radiomics for pre-intervention clot phenotyping, highlighting its potential to inform future stroke treatment planning.

## MATERIALS AND METHODS

### Sample Preparation

We obtained fresh human whole blood from 5 healthy adult volunteers (ages 21-26 years; 2 men and 3 women) under an institutionally approved protocol. All subjects provided informed consent. We collected blood intended for clot formation into acid citrate dextrose (ACD) anticoagulant. Separately, we collected a small aliquot from each volunteer into ethylenediaminetetraacetic acid (EDTA) anticoagulant and submitted it to a local clinical laboratory for complete blood count testing to determine baseline hematocrit.

We split blood into 2 aliquots. One aliquot was centrifuged at 3200 RPM for 10 minutes at room temperature to separate plasma and red blood cells (RBCs)^24^. We then recombined plasma and RBCs to generate clots with target RBC volume fractions (hematocrits) of 0%, 10%, 20%, 60%, and 80%. The second aliquot was used to form native whole blood clots. To initiate coagulation, we added calcium chloride to all samples to a final concentration of 20 mM^25–27^. For both the recombined clots and the native whole blood clots, we prepared 2 conditions: one without thrombin and one with thrombin (1 U/mL).

Clots coagulated in the presence of thrombin consistently underwent volumetric contraction. We refer to these as the thrombin (*contracted*) condition. Clots coagulated without thrombin showed minimal contraction and are referred to as the no thrombin (*uncontracted*) condition (Fig 1). Throughout the manuscript, hematocrit values are defined by the source blood composition used to prepare each clot. In secondary analyses, we estimated an effective post-contraction hematocrit by normalizing the initial hematocrit by the measured volumetric contraction. For this calculation, we assumed that the total RBC number was conserved within the clot during contraction (Supplemental Data Figs 2, 5-6, Table 3).

**Fig 1.**
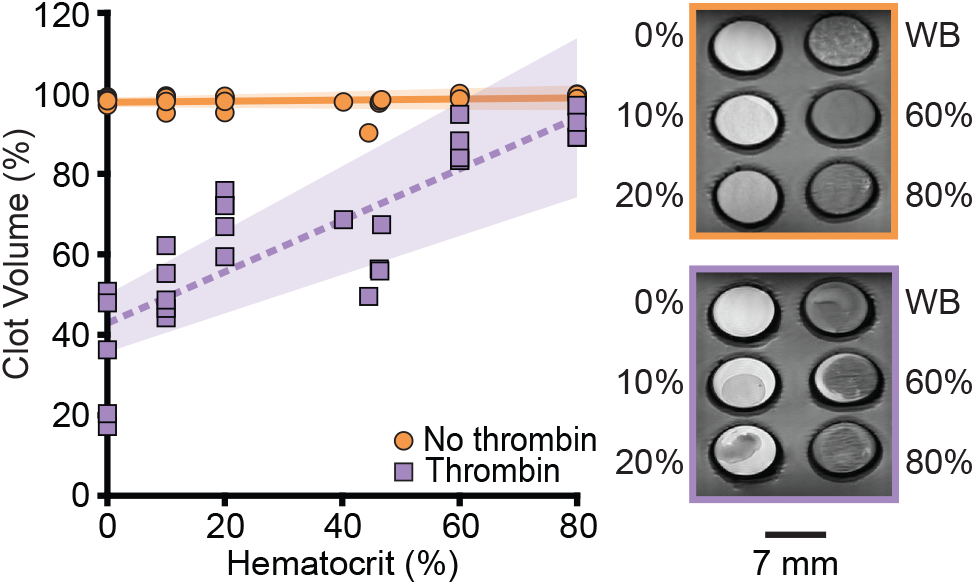
Relationship between clot contraction and hematocrit. Representative T2-weighted MR images show clots coagulated with and without thrombin. The scatterplot shows clot volume (expressed as a percentage of the initial volume) versus hematocrit for individual blood clots. Each data point represents a single clot, and they are colored by condition: round orange indicates clots coagulated without thrombin, and square purple indicates clots coagulated with thrombin. Clots without thrombin show no relationship between contraction and hematocrit (R^2^ = 0.04, p = 0.279), whereas clots with thrombin show a strong positive relationship (R^2^ = 0.71, p < 0.001). Linear regression fits are shown for each condition (solid orange, dashed purple) with shaded 95% CIs. WB denotes whole blood.

We incubated clots for 2 hours to allow coagulation and then embedded them in an agarose-gelatin-glycerol hydrogel to maintain sample position during MRI.

### Image Acquisition

We acquired all images using a 7 T preclinical MRI scanner (Bruker Biospec, Billerica, MA, USA) equipped with a 38 mm single-channel volume coil. We included quantitative mapping sequences, clinically established stroke MRI sequences, and investigational sequences not previously used in stroke imaging. For all scans, the slice thickness was 1 mm, and the field of view was 40 x 32 mm.

Quantitative imaging included T2 mapping with a multi-echo spin-echo sequence, T1 mapping with a variable flip angle gradient-recalled echo sequence, and diffusion-weighted imaging for apparent diffusion coefficient estimation. Clinical sequences included T2-weighted rapid acquisition with relaxation enhancement (RARE), SWI, and T1-weighted GRE imaging. Investigational sequences included flow-sensitive alternating inversion recovery (FAIR) RARE, ultra-short echo time (UTE), and T1-weighted FLASH imaging. In the absence of flow, FAIR RARE produces a relaxation-weighted signal dominated by inversion recovery and T2-weighted fast spin-echo signal evolution. Detailed sequence parameters are provided in Supplemental Data Table 1.

### Image Processing

For each subject and MRI scan, we loaded reconstructed images into MATLAB (Version R2024a; MathWorks, Natick, MA, USA) and scaled them using the Bruker reconstruction factor. For quantitative images, we generated voxel-wise parametric maps. To do so, we fit signal decay across echo times to a mono-exponential model to estimate T2, used a variable flip angle model of spoiled gradient-echo signal to estimate T1, and applied log-linear regression of diffusion-weighted signal versus b-value to estimate the ADC. Nonphysical voxel estimates were excluded from fits. For qualitative images (e.g., standard magnitude images), we applied z-score normalization.

For each scan, we selected three central slices and manually delineated six circular regions of interest (ROIs) corresponding to the clot conditions. Each ROI had a fixed diameter of 4 mm. We then calculated the average quantitative values (T1, T2, and ADC) or the average signal intensity (qualitative images) across the three slices for each clot.

### Radiomics

Radiomic analysis was performed separately for each MRI scan type. Prior to feature extraction, we resampled images to an isotropic in-plane resolution defined by the smallest native pixel dimension (see Supplemental Data Table 1). Linear interpolation was applied to image data and nearest-neighbor interpolation was applied to ROI masks. We then extracted 2D radiomic features from each ROI using PyRadiomics (v2.2) with 40 intensity bins per image^28^. Features included first-order, shape, and texture measures^29^. Averaging these features across the three slices yielded a single value per feature for each clot and scan.

### Statistical Analysis

We conducted all analyses in MATLAB. We analyzed clots coagulated with and without thrombin separately.

#### Signal-Hematocrit Correlations

To assess associations between MRI-derived measurements and clot composition, we correlated quantitative parameters (T1, T2, ADC) and qualitative image signal intensities with hematocrit. We evaluated linear relationships using ordinary least squares regression and reported R^2^ and p-values for each association.

#### Radiomic Feature Regression with Clot Hematocrit

To evaluate the relationship between radiomic features and clot hematocrit, we performed regression analysis for each clot condition and scan type. Analyses were also conducted using pooled features across scans. To capture linear effects and potential nonlinear and higher-order relationships, we used three modeling approaches: standard least absolute shrinkage and selection operator (LASSO) regression, polynomial LASSO regression, and random forest regression. Polynomial LASSO was implemented using standard MATLAB feature expansion to include higher-order terms prior to LASSO regularization.

Data separation and feature preprocessing were identical across regression models. Specifically, we used 5-fold subject-level cross-validation to ensure independence between training and test sets. Within each training set, we removed low-variance and highly correlated features and standardized the remaining features. Shape features were excluded because ROIs had identical geometry. Identical preprocessing steps were applied to the held-out test fold.

Model performance was quantified solely from cross-validated predictions on held-out subjects. We assessed performance using R^2^ and root-mean-square error (RMSE). Negative predictions were clipped to zero.

#### Classification of Clot Composition and Contraction State

To test whether radiomic features could accurately distinguish clot types, we trained random forest classifiers on two tasks. The first task discriminated between RBC-rich clots (native whole blood, 60%, 80% hematocrit) and fibrin-rich clots (0%, 10%, 20% hematocrit). The second task discriminated between clots coagulated with thrombin (contracted) and without thrombin (uncontracted). For composition, we performed classification separately for clots coagulated with and without thrombin. Furthermore, analyses were carried out for each MRI scan type and using pooled radiomic features.

Feature preprocessing and model training mirrored the steps applied in regression analyses. We used 5-fold subject-level cross-validation, excluded shape features, and optimized model hyperparameters via inner cross-validation. We assessed classifier performance using the area under the receiver operating characteristic curve (AUC) and 95% CIs, calculated from cross-validated predictions.

## RESULTS

### Correlation of Quantitative MRI Parameters and Signal Intensity with Clot Hematocrit

Measuring clot hematocrit prior to intervention would be clinically useful. To this end, we compared nine MRI sequences spanning quantitative, clinical, and investigational approaches. Across these sequences, hematocrit showed consistent associations with MRI signal and quantitative parameters, although the strength of these relationships varied by sequence type (Fig 2, Table 1).

**Table 1.** Regression metrics of MRI measurements versus clot hematocrit.

| Sequence | Condition | R | R<sup>2</sup> | p-value |
| --- | --- | --- | --- | --- |
| T2 Mapping | No thrombin | -0.86 | 0.72 | <0.001 |
|  | Thrombin | -0.75 | 0.56 | <0.001 |
| T1 Mapping | No thrombin | -0.82 | 0.68 | <0.001 |
|  | Thrombin | -0.80 | 0.64 | <0.001 |
| ADC Mapping | No thrombin | -0.92 | 0.85 | <0.001 |
|  | Thrombin | -0.90 | 0.81 | <0.001 |
| T2 RARE | No thrombin | -0.79 | 0.62 | <0.001 |
|  | Thrombin | -0.71 | 0.51 | <0.001 |
| SWI | No thrombin | -0.79 | 0.62 | <0.001 |
|  | Thrombin | -0.71 | 0.50 | <0.001 |
| T1 GRE | No thrombin | -0.68 | 0.46 | <0.001 |
|  | Thrombin | -0.68 | 0.46 | <0.001 |
| FAIR RARE | No thrombin | -0.84 | 0.71 | <0.001 |
|  | Thrombin | -0.77 | 0.60 | <0.001 |
| UTE | No thrombin | -0.21 | 0.04 | 0.304 |
|  | Thrombin | -0.29 | 0.08 | 0.127 |
| T1 FLASH | No thrombin | -0.33 | 0.11 | 0.071 |
|  | Thrombin | -0.39 | 0.15 | 0.034 |

**Fig 2.**
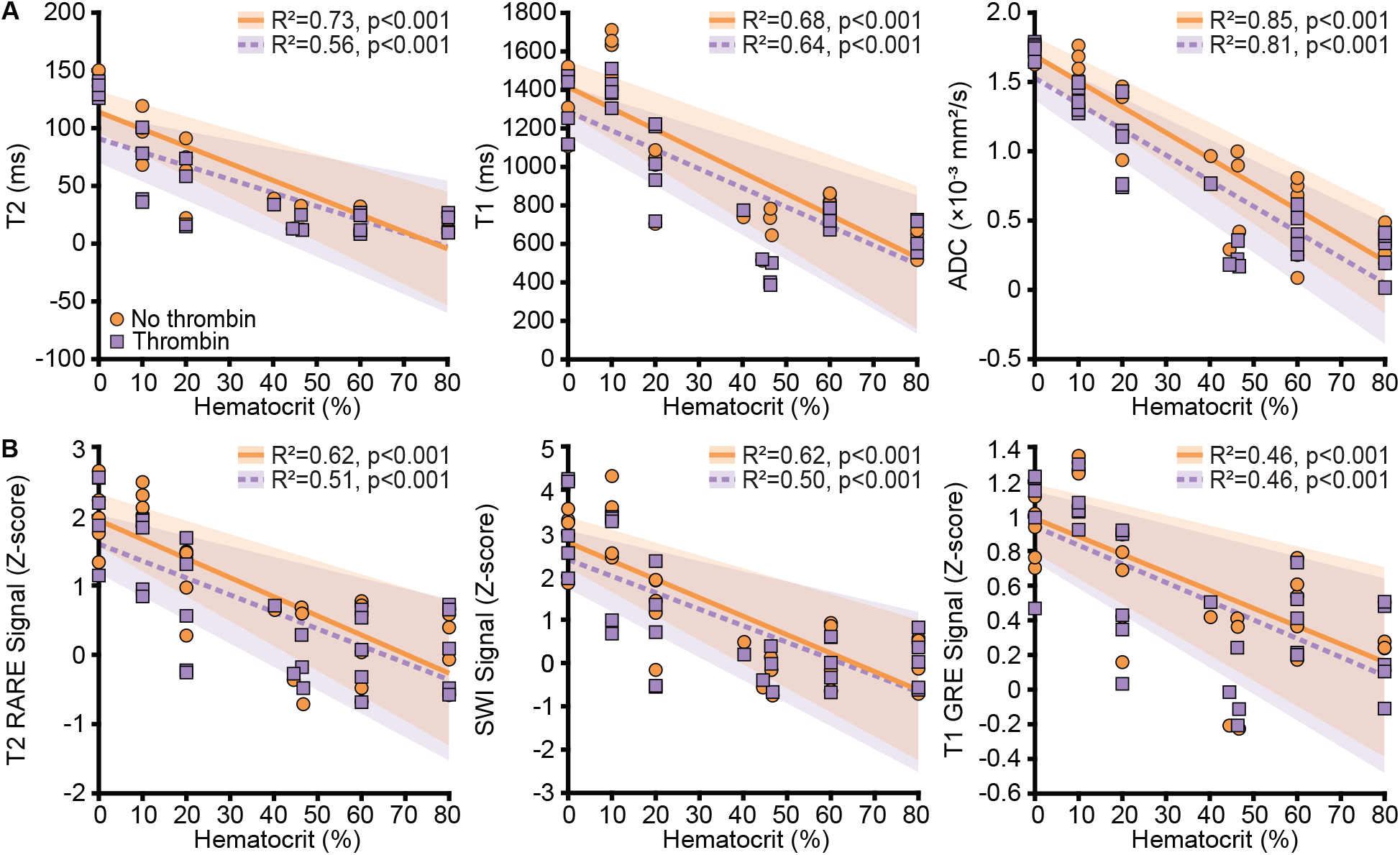
Clot hematocrit is associated with quantitative MRI parameters and signal intensity across sequences and conditions. Scatterplots show (A) T1, T2, and ADC versus hematocrit and (B) T2 RARE, SWI, and T1 GRE signal intensity versus hematocrit. Each data point represents a single clot. Data points are colored by condition: round orange indicates clots coagulated without thrombin, and square purple indicates clots coagulated with thrombin. Linear regression fits are shown for each condition (solid orange, dashed purple) with shaded 95% CIs.

Quantitative sequences were the best overall: increasing hematocrit was associated with decreases in T1, T2, and ADC. In fact, ADC showed the strongest relationship with hematocrit among all sequences (Fig 2A). Clinical sequences followed similar trends but with weaker associations (Fig 2B). Among these, T2 RARE and SWI outperformed T1 GRE. In contrast, investigational sequences showed more variable behavior (Supplemental Data Fig 1). UTE and T1 FLASH performed the worst, showing weak or insignificant associations with hematocrit. Conversely, FAIR RARE outperformed all clinical sequences, showing associations comparable to T1 and T2.

Across all sequences with a significant relationship with hematocrit, correlations were comparable or weaker in thrombin-treated clots. Using post-contraction corrected hematocrit values did not change these overall patterns (Supplemental Table S1).

### Radiomic Feature-Based Prediction of Clot Hematocrit

Having confirmed that hematocrit is associated with MRI signal across sequences, we next tested whether radiomic features could extract additional predictive information from these images. We tested multiple modeling approaches to account for different patterns of feature interaction, and random forest consistently provided the strongest predictions. Accordingly, Fig 3 and Table 2 summarize random forest results. Comparative metrics for all models are in Supplemental Data Table 2.

**Table 2.** Random forest regression metrics of radiomic features versus clot hematocrit.

| Sequence | Condition | R | $R^2$ | RMSE |
| --- | --- | --- | --- | --- |
| T2 Mapping | No thrombin | 0.85 | 0.79 | 0.16 |
|  | Thrombin | 0.76 | 0.56 | 0.19 |
| T1 Mapping | No thrombin | 0.94 | 0.87 | 0.10 |
|  | Thrombin | 0.88 | 0.76 | 0.14 |
| ADC Mapping | No thrombin | 0.90 | 0.81 | 0.12 |
|  | Thrombin | 0.89 | 0.79 | 0.13 |
| T2 RARE | No thrombin | 0.81 | 0.66 | 0.17 |
|  | Thrombin | 0.74 | 0.52 | 0.19 |
| SWI | No thrombin | 0.79 | 0.62 | 0.17 |
|  | Thrombin | 0.57 | 0.31 | 0.24 |
| T1 GRE | No thrombin | 0.79 | 0.61 | 0.12 |
|  | Thrombin | 0.69 | 0.47 | 0.21 |
| FAIR RARE | No thrombin | 0.85 | 0.72 | 0.15 |
|  | Thrombin | 0.66 | 0.43 | 0.21 |
| UTE | No thrombin | -0.40 | -0.23 | 0.28 |
|  | Thrombin | -0.20 | -0.16 | 0.30 |
| T1 FLASH | No thrombin | -0.18 | -0.28 | 0.17 |
|  | Thrombin | 0.58 | 0.33 | 0.23 |
| Pooled Features | No thrombin | 0.89 | 0.75 | 0.14 |
|  | Thrombin | 0.90 | 0.78 | 0.13 |

**Table 3.**
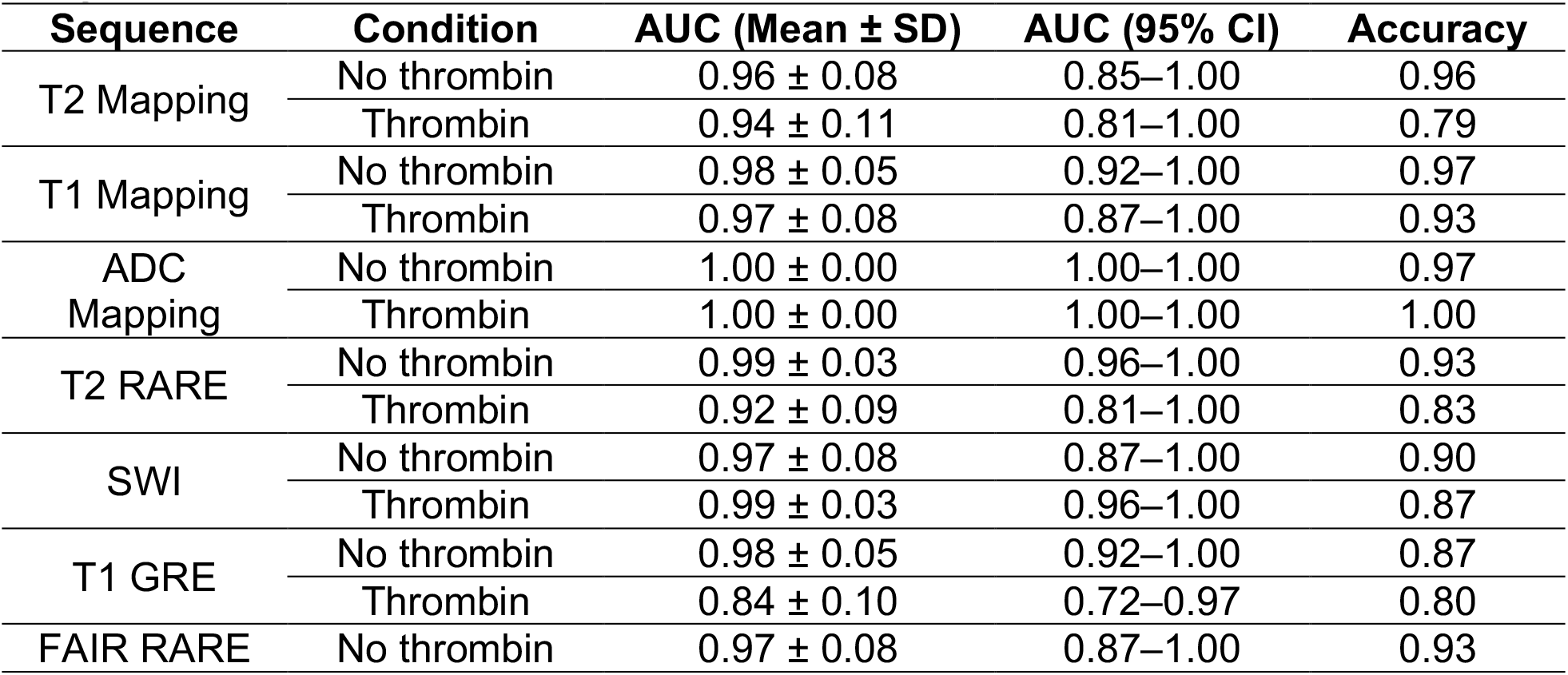
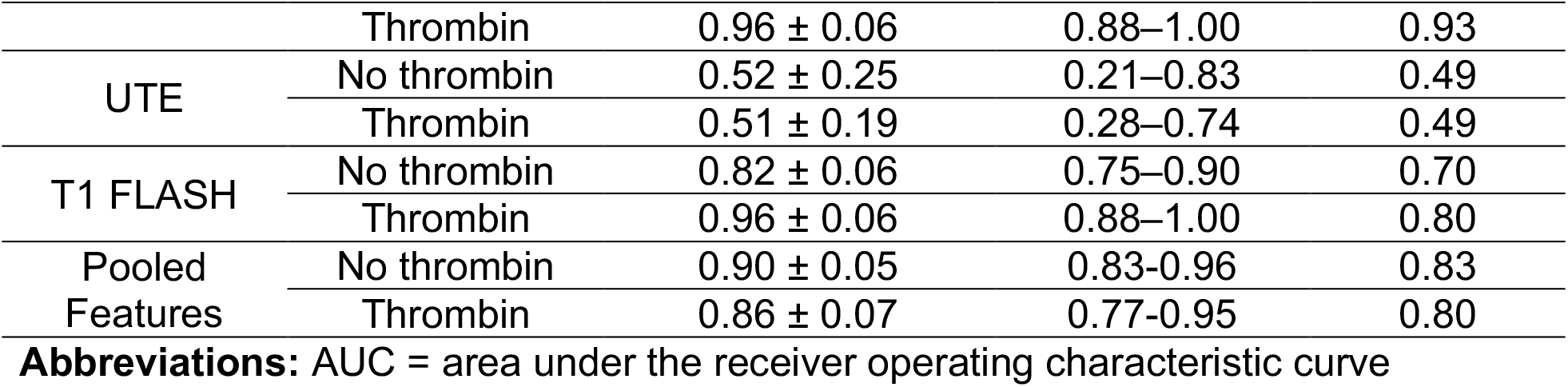
Classification performance for discrimination of RBC-rich and fibrin-rich clots using radiomic features from individual MRI sequences and pooled features.

**Fig 3.**
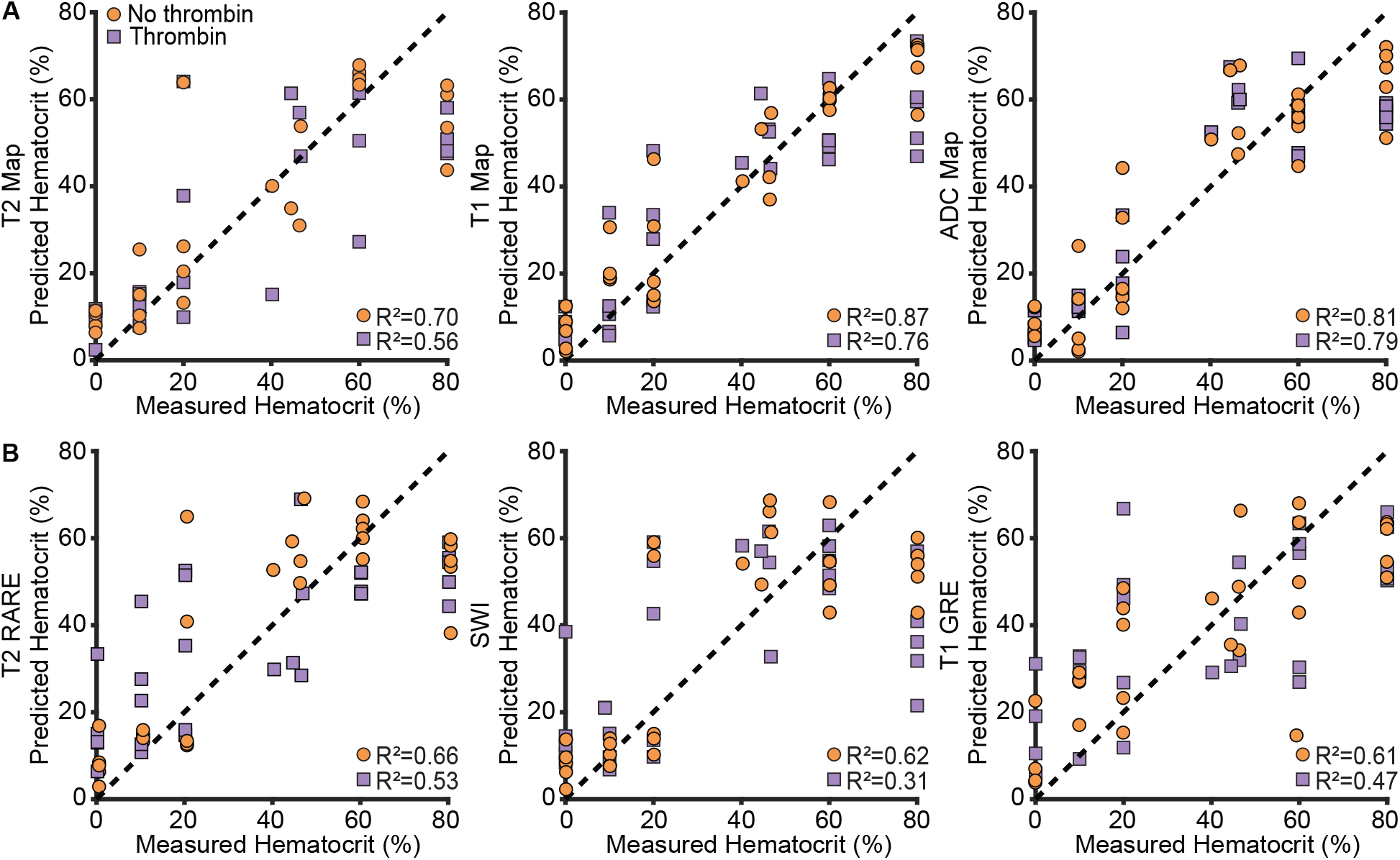
Random forest regression shows sequence-dependent variability in predicting clot hematocrit from radiomic features. (A) Predictions based on radiomic features extracted from T2, T1, and ADC maps. (B) Predictions based on radiomic features from T2 RARE, SWI, and T1 GRE images. Each data point represents a single clot. Data points are colored by condition: round orange indicates clots coagulated without thrombin, and square purple indicates clots coagulated with thrombin. The dotted line indicates perfect prediction (y = x). Regressions are based on 5-fold cross-validation.

Compared with quantitative parameters and signal intensity, radiomic features improved hematocrit prediction only for some sequences. Among all sequences, quantitative scans again provided the strongest predictions (Fig 3A). However, in this case, T1 maps were the best overall, followed closely by ADC and T2 maps. Likewise, clinical sequences offered moderately strong predictions (Fig 3B). Among these, T2 RARE outperformed both SWI and T1 GRE. Investigational sequences, by contrast, had the weakest predictions overall (Supplemental Data Fig 3). While FAIR RARE achieved predictions comparable to the clinical sequences, UTE and T1 FLASH yielded weak predictions.

Across sequence types, predictions were slightly weaker in thrombin-treated clots. Furthermore, pooled feature models strongly predicted hematocrit, with performance comparable to the best individual quantitative sequences (Supplemental Data Fig 4). Finally, using post-contraction corrected hematocrit values did not change these overall patterns (Supplemental Data Figs 5–6, Table 3).

### Radiomic Feature-Based Classification of RBC- and Fibrin-Rich Clots

After finding that radiomic features correlate with clot hematocrit, we next tested whether they could distinguish RBC-rich from fibrin-rich clots. Importantly, this distinction may be sufficient to guide treatment, and eight of our nine sequences enabled very strong discrimination (Table 3, Fig 4).

**Fig 4.**
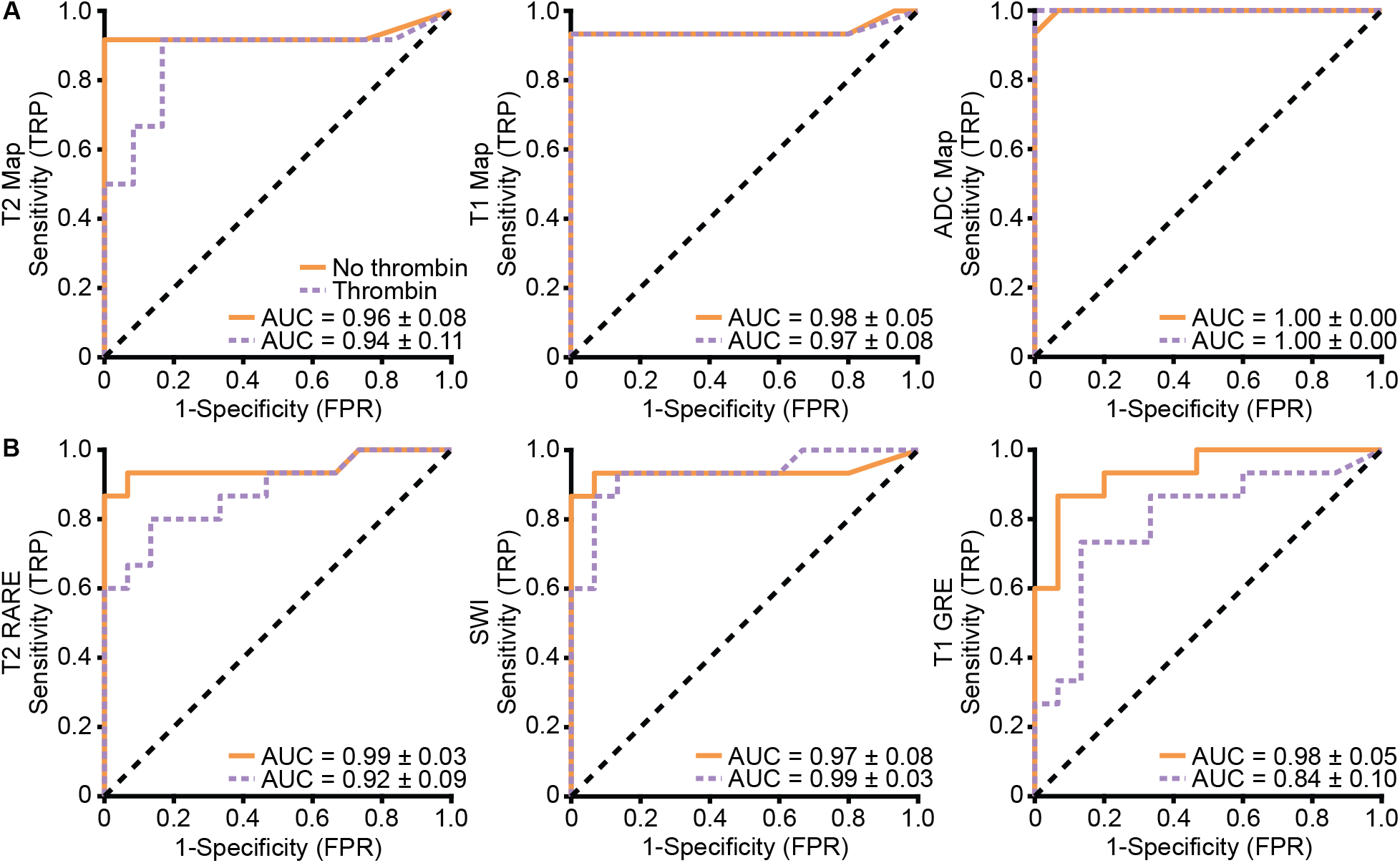
ROC analysis shows strong performance of radiomic features in discriminating RBC-rich versus fibrin-rich clots across MRI sequences. (A) Classifications based on radiomic features extracted from T2, T1, and ADC maps. (B) Classifications based on radiomic features extracted from T2 RARE, SWI, and T1 GRE scans. Each curve represents the mean ROC across 5-fold cross-validation. Curves are colored by condition: orange indicates clots coagulated without thrombin, and purple indicates clots coagulated with thrombin. The area under the curve (AUC) ± standard deviation is shown on the plots.

ADC features stood out: they achieved near-perfect classification (AUC = 1.00). The remaining quantitative sequences, along with all clinical sequences, demonstrated excellent discrimination (AUCs ≥ 0.84). In contrast, investigational sequences were more variable (AUC range: 0.51–0.97). Interestingly, T1 FLASH, despite weak performance in regression, showed strong classification. UTE, on the other hand, yielded near-chance discrimination (Supplemental Data Fig 7).

For most sequence types, discrimination was slightly weaker in thrombin-treated clots. Additionally, combining features across all MRI sequences yielded strong classification, although performance was similar to the individual quantitative or clinical sequences (Supplemental Data Fig 8).

### Radiomic Feature-Based Classification of Uncontracted and Contracted Clots

Because contraction increases clot density, detecting it could help guide treatment. We therefore tested whether radiomic features could distinguish clots coagulated with thrombin (contracted) from those coagulated without thrombin (uncontracted). Overall, discrimination was more variable than for hematocrit-based classification (Table 4, Fig 5, Supplemental Data Fig 9).

**Table 4.**
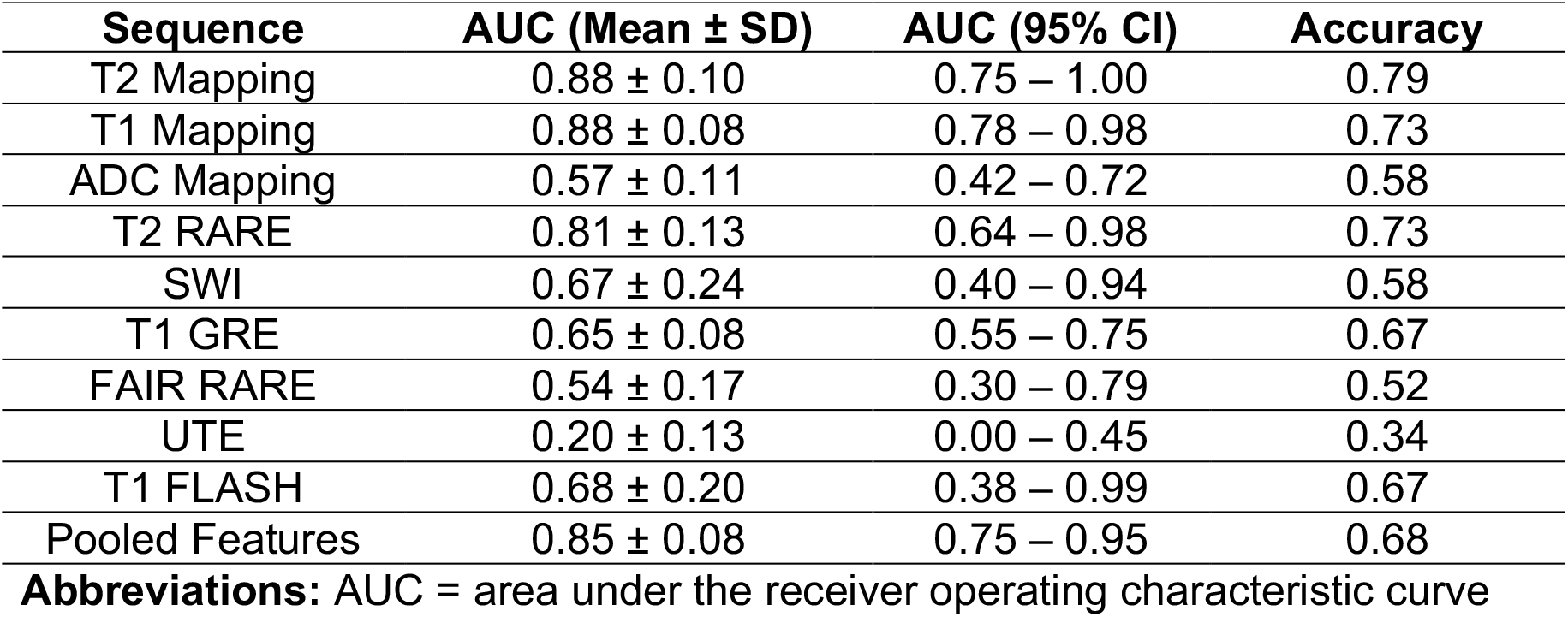
Classification performance for discrimination of clots coagulated with thrombin (contracted) versus without thrombin (uncontracted) using radiomic features from individual MRI sequences and pooled features.

| Sequence | AUC (Mean ± SD) | AUC (95% CI) | Accuracy |
| --- | --- | --- | --- |
| T2 Mapping | $0.88 \pm 0.10$ | 0.75 – 1.00 | 0.79 |
| T1 Mapping | $0.88 \pm 0.08$ | 0.78 – 0.98 | 0.73 |
| ADC Mapping | $0.57 \pm 0.11$ | 0.42 – 0.72 | 0.58 |
| T2 RARE | $0.81 \pm 0.13$ | 0.64 – 0.98 | 0.73 |
| SWI | $0.67 \pm 0.24$ | 0.40 – 0.94 | 0.58 |
| T1 GRE | $0.65 \pm 0.08$ | 0.55 – 0.75 | 0.67 |
| FAIR RARE | $0.54 \pm 0.17$ | 0.30 – 0.79 | 0.52 |
| UTE | $0.20 \pm 0.13$ | 0.00 – 0.45 | 0.34 |
| T1 FLASH | $0.68 \pm 0.20$ | 0.38 – 0.99 | 0.67 |
| Pooled Features | $0.85 \pm 0.08$ | 0.75 – 0.95 | 0.68 |

**Abbreviations:** AUC = area under the receiver operating characteristic curve

**Fig 5.**
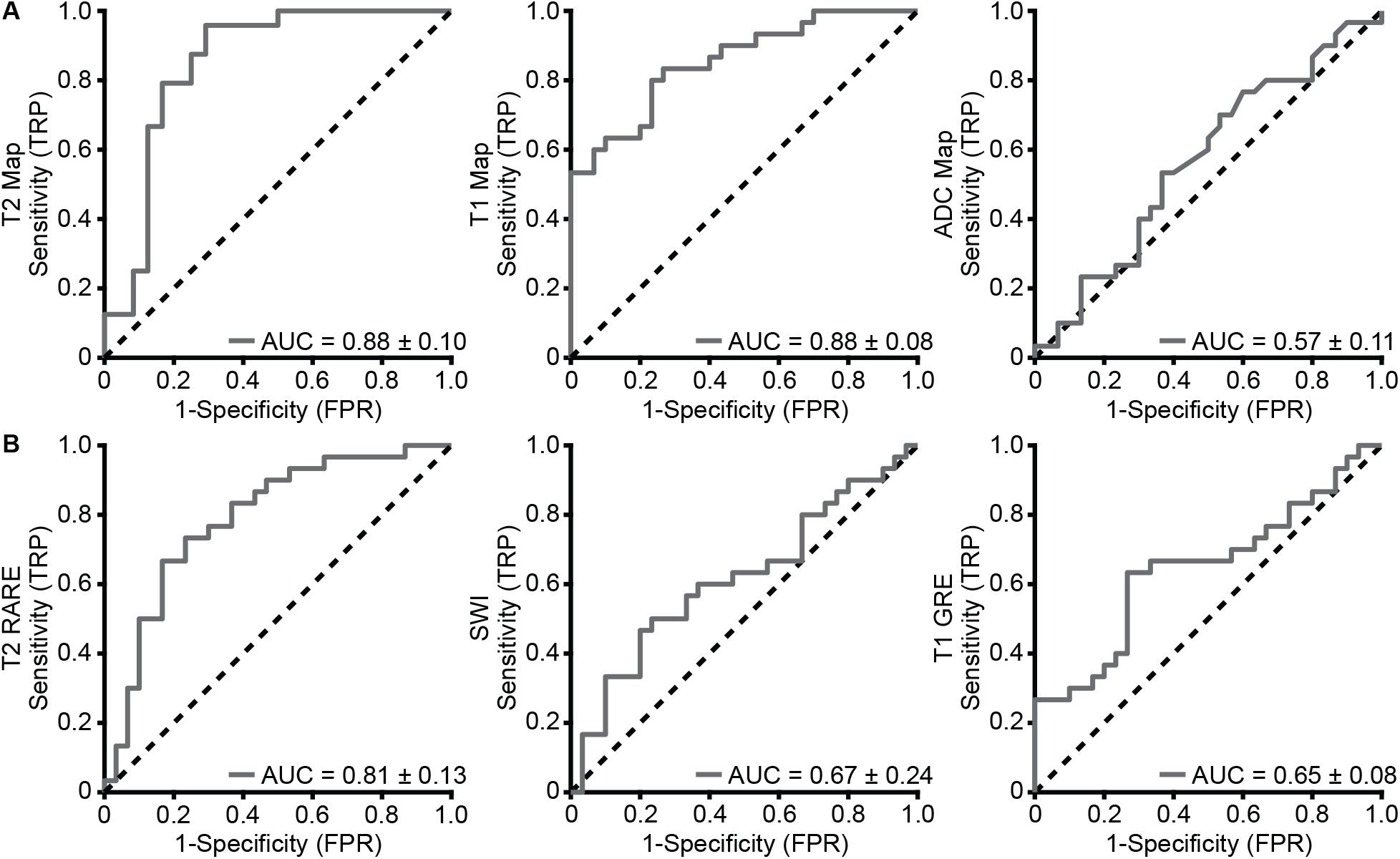
ROC analysis shows variable performance of radiomic features in discriminating clots coagulated with thrombin (contracted) versus without thrombin (uncontracted). (A) Classifications based on radiomic features extracted from T2, T1, and ADC maps. (B) Classifications based on radiomic features extracted from T2 RARE, SWI, and T1 GRE scans. Each curve represents the mean ROC across 5-fold cross-validation. The area under the curve (AUC) ± standard deviation is shown on the plots.

Among all sequences, T1 maps, T2 maps, and T2 RARE were the only scans with strong discriminatory power (AUCs ≥ 0.81). All other sequences demonstrated poor discriminatory power (AUC range: 0.20–0.68), including ADC maps, which excelled for hematocrit prediction and classification. Combining features across all sequences yielded strong classification, comparable to T1 and T2 maps and T2 RARE alone (Supplemental Data Fig 10).

## DISCUSSION

Our goal was to test whether MRI combined with radiomic analysis can predict clot composition and contraction in vitro. To this end, we evaluated how MRI signal intensities, quantitative parameters, and radiomic features relate to clot hematocrit and enable classification by composition and contraction state. We found that MRI-based radiomic analysis can estimate clot hematocrit and differentiate clots by composition and, to a lesser extent, contraction. Quantitative MRI sequences demonstrated strong performance across both regression and classification tasks, while clinical stroke sequences also showed strong discriminative performance for clot composition.

First, our findings show that MRI signal and quantitative parameters are strongly associated with clot hematocrit. These results are consistent with prior findings in clot analogs and retrieved thrombi, which have demonstrated that T1, T2, and T2* decrease with increasing clot hematocrit^15,30–34^. In fact, prior work has shown that clinical sequences, such as susceptibility-weighted techniques, are sensitive to clot composition^32,35^. However, sensitivity was limited to intermediate hematocrit ranges^35^. In contrast, our study demonstrates consistent signal changes across a broad range of hematocrits and across multiple MRI approaches, including quantitative, clinical, and investigational sequences. Although some sequences showed signal saturation at higher hematocrits, overall trends remained well-defined. Thus, our study demonstrates that both quantitative MRI techniques and clinical stroke protocols capture clot compositional changes. However, the extent to which these relationships translate to in vivo imaging remains unclear. In vivo, MRI markers of composition are limited, with the most well-established being the susceptibility-related blooming artifact. This hypointense signal surrounding the occluding clot is associated with RBC-rich composition^10,36^. The signal loss we observe at higher hematocrits is consistent with this imaging feature. However, beyond the blooming artifact, many of the signal relationships identified in this study are not readily observed in standard clinical imaging^37^. This likely reflects our simplified in vitro setting, in which clots are imaged in isolation without surrounding brain tissue or vasculature. Thus, subtle compositional effects may be obscured in vivo, underscoring the need for more sensitive analytical approaches. In this regard, radiomic analysis enables the capture and quantification of image features and spatial heterogeneity that are not visually apparent.

Second, our data show that radiomic features improved hematocrit prediction for some sequences, but not across all scans. In contrast, radiomics enabled consistently strong discrimination between RBC- and fibrin-rich clots. Notably, we observed strong classification with radiomic features from both quantitative MRI and clinically used stroke sequences. This suggests that existing stroke protocols may be able to distinguish clot composition in vivo, which could ultimately inform treatment decisions. However, MRI-based radiomic studies for clot phenotyping remain limited. Prior work has primarily focused on radiomic analysis of DWI, where features have been shown to predict long-term outcomes and overall prognosis^38,39^. In contrast, CT-based radiomic studies are more extensive and have demonstrated the ability to predict thrombectomy outcomes, infer stroke etiology, and estimate clot composition^21–23,40–44^. Because these clinical endpoints are influenced by clot composition, our findings suggest that MRI-based radiomic analysis could provide similar clinical value. Additionally, MRI offers multiple contrasts that may capture complementary clot properties, potentially providing more detailed insight than CT alone.

Finally, we found that only radiomic features from select sequences could discriminate clots based on contraction. Specifically, only features from T1 and T2 maps and T2 RARE showed effective classification. Our findings are consistent with prior work demonstrating that T2-weighted spin echo and quantitative T2 measurements are sensitive to clot contraction, although these studies did not incorporate radiomic analysis^30,45^. Interestingly, features from ADC mapping did not demonstrate strong discriminatory power, despite contraction being known to decrease clot diffusivity^8,46^. This difference may reflect a greater sensitivity of relaxation-based MRI sequences to changes in clot density and structure. However, further studies are needed to better understand how contraction-related microstructural changes are captured by different MRI parameters.

Our study has important limitations. First, we formed homogeneous clots in vitro under controlled conditions and without flow. This approach was designed to create a reproducible model for studying the relationship between MRI signal and composition. However, in vivo clots are heterogeneous and shaped by hemodynamics, which will affect MRI signal^5,47^. Thus, future studies should extend our approach to heterogeneous clots that better capture in vivo compositional and structural variability. Second, we performed imaging using a 7 T scanner. This provides a higher signal-to-noise ratio and spatial resolution than typical clinical 1.5-3 T scanners that are used clinically^48^. Similarly, clots were imaged in isolation, without surrounding brain tissue or vasculature. Both factors likely enhanced our ability to detect and differentiate MRI features. Thus, predictive performance would likely be reduced in vivo. Future work should therefore evaluate our findings at clinical field strengths under more realistic imaging conditions. Third, our sample size was relatively small. Nevertheless, the study was sufficiently powered to detect patterns in correlations, regression outcomes, and classification performance across sequences.

## CONCLUSION

MRI-based radiomic analysis can characterize clot composition and contraction in vitro. Notably, sequences already used in clinical stroke protocols effectively distinguished RBC-rich from fibrin-rich clots, with some showing sensitivity to contraction. Our findings motivate further validation in clinically relevant imaging conditions and in vivo studies.

## ACKNOWLEDGEMENTS

This work was supported by the National Science Foundation through grant 2046148. We acknowledge the use of the Biomedical Imaging Center (BIC) at The University of Texas at Austin (RRID: SCR_021728), whose facilities and resources contributed to this research.

