## Supplemental Data for "MRI-Based Blood Clot Phenotyping: An In Vitro Study"

### SUPPLEMENTARY INFORMATION

**Table S1.** MRI sequence parameters.

| Sequence | TE (ms) | TR (ms) | Flip Angle (°) | Other Parameters | In-plane Resolution (mm) |
|---|---|---|---|---|---|
| T2 Mapping | 6.66–73.27 (11 echoes) | 8799 | — | Multi-echo spin-echo | 0.2222 × 0.266 |
| T1 Mapping | — | 200 | 5, 10, 15, 20, 45 | Variable flip angle GRE | 0.3125 × 0.5 |
| DWI (ADC Mapping) | 28.7 | 2500 | — | b = 150, 300, 800 s/mm² (single direction) | 0.3125 × 0.25 |
| T2 RARE | 14 | 3500 | — | RARE factor = 8 | 0.1563 × 0.125 |
| SWI | 12 | 722.75 | 40 | Single-echo GRE | 0.1042 × 0.0833 |
| T1 GRE | 2.65 | 494 | 30 | Spoiled GRE | 0.3125 × 0.25 |
| FAIR RARE | 80 (effective) | 4309 | — | TI = 1000 ms; echo spacing = 5 ms; RARE factor = 32 | 0.3125 × 0.25 |
| UTE | 0.8 | 10 | 15 | — | 0.3125 × 0.25 |
| T1 FLASH | 2.71 | 150 | 24 | Spoiled GRE | 0.156 × 0.125 |

**Abbreviations:** RARE = rapid acquisition with relaxation enhancement; FAIR = flow-sensitive alternating inversion recovery; UTE = ultrashort echo time

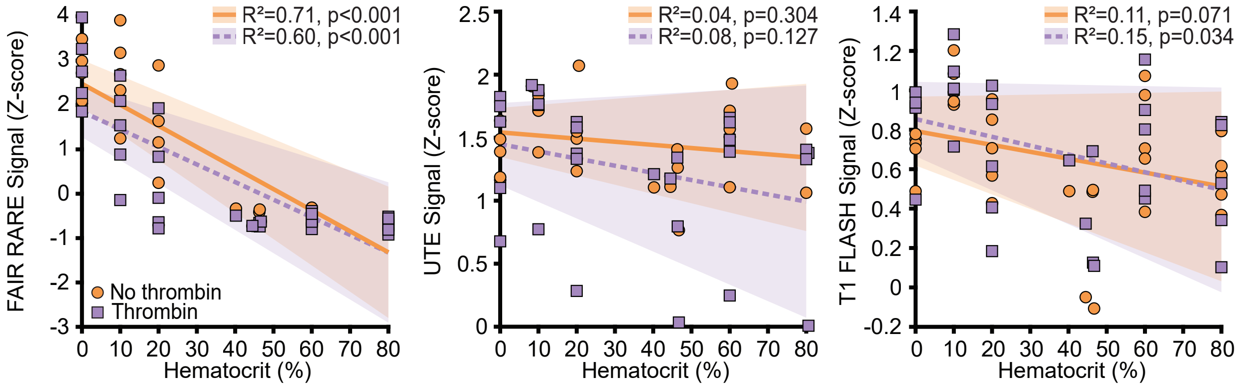


**Fig S1.** Clot hematocrit shows sequence-dependent variation in its association with signal intensity across FAIR RARE, UTE, and T1 FLASH sequences. Each data point represents a single clot. Data points are colored by condition: round orange indicates clots coagulated without thrombin, and square purple indicates clots coagulated with thrombin. Linear regression fits are shown for each condition (solid orange, dashed purple) with shaded 95% CIs.

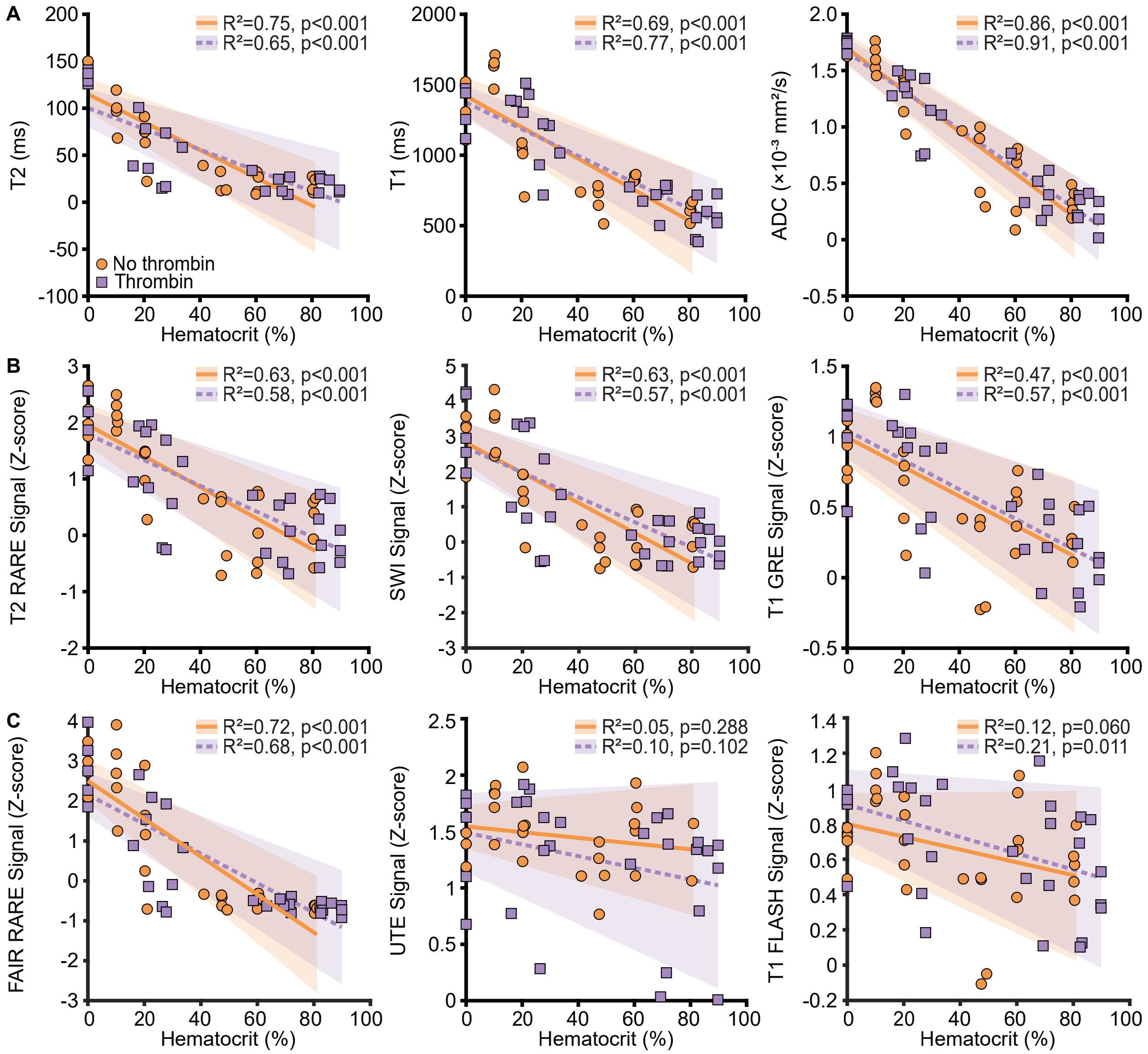


**Fig S2.** Corrected clot hematocrit is associated with quantitative MRI parameters and signal intensity from some sequences. Scatterplots show (A) T1, T2, and ADC versus corrected hematocrit, (B) T2 RARE, SWI, and T1 GRE signal intensity versus corrected hematocrit, and (C) FAIR RARE, UTE, and T1 FLASH signal intensity versus corrected hematocrit. Each data point represents a single clot. Data points are colored by condition: round orange indicates clots coagulated without thrombin, and square purple indicates clots coagulated with thrombin. Linear regression fits are shown for each condition (solid orange, dashed purple) with shaded 95% CIs.

**Table S2.** Performance metrics for radiomic feature regression models predicting hematocrit from individual MRI sequences and pooled features.

| Scan | Condition | Model | R | $R^2$ | RMSE |
|---|---|---|---|---|---|
| T2 Mapping | No thrombin | Linear LASSO | 0.81 | 0.57 | 0.19 |
| | | Poly LASSO | 0.66 | 0.42 | 0.22 |
| | | Random Forest | 0.84 | 0.70 | 0.16 |
| | Thrombin | Linear LASSO | 0.64 | 0.41 | 0.22 |
| | | Poly LASSO | 0.55 | 0.27 | 0.24 |
| | | Random Forest | 0.76 | 0.56 | 0.19 |
| T1 Mapping | No thrombin | Linear LASSO | 0.90 | 0.77 | 0.14 |
| | | Poly LASSO | 0.89 | 0.77 | 0.14 |
| | | Random Forest | 0.94 | 0.87 | 0.10 |
| | Thrombin | Linear LASSO | 0.73 | 0.52 | 0.20 |
| | | Poly LASSO | 0.76 | 0.56 | 0.19 |
| | | Random Forest | 0.88 | 0.76 | 0.14 |
| ADC Mapping | No thrombin | Linear LASSO | 0.20 | 0.03 | 0.28 |
| | | Poly LASSO | 0.33 | 0.11 | 0.27 |
| | | Random Forest | 0.90 | 0.81 | 0.12 |
| | Thrombin | Linear LASSO | 0.38 | 0.11 | 0.27 |
| | | Poly LASSO | 0.47 | 0.07 | 0.27 |
| | | Random Forest | 0.89 | 0.79 | 0.13 |
| T2 RARE | No thrombin | Linear LASSO | 0.72 | 0.50 | 0.20 |
| | | Poly LASSO | 0.60 | 0.31 | 0.24 |
| | | Random Forest | 0.81 | 0.66 | 0.17 |
| | Thrombin | Linear LASSO | 0.84 | 0.63 | 0.17 |
| | | Poly LASSO | 0.78 | 0.58 | 0.18 |
| | | Random Forest | 0.74 | 0.53 | 0.19 |
| SWI | No thrombin | Linear LASSO | 0.73 | 0.50 | 0.20 |
| | | Poly LASSO | 0.77 | 0.53 | 0.19 |
| | | Random Forest | 0.79 | 0.62 | 0.17 |
| | Thrombin | Linear LASSO | 0.69 | 0.46 | 0.21 |
| | | Poly LASSO | 0.70 | 0.46 | 0.21 |
| | | Random Forest | 0.57 | 0.31 | 0.24 |
| T1 GRE | No thrombin | Linear LASSO | 0.50 | 0.08 | 0.27 |
| | | Poly LASSO | 0.92 | 0.81 | 0.12 |
| | | Random Forest | 0.79 | 0.61 | 0.18 |
| | Thrombin | Linear LASSO | 0.70 | 0.41 | 0.22 |
| | | Poly LASSO | 0.68 | 0.44 | 0.21 |
| | | Random Forest | 0.69 | 0.47 | 0.21 |
| FAIR RARE | No thrombin | Linear LASSO | 0.81 | 0.59 | 0.18 |
| | | Poly LASSO | 0.76 | 0.57 | 0.19 |
| | | Random Forest | 0.85 | 0.72 | 0.15 |
| | Thrombin | Linear LASSO | 0.72 | 0.45 | 0.21 |
| | | Poly LASSO | 0.63 | 0.38 | 0.22 |

| | | | | | |
|---|---|---|---|---|---|
| | | Random Forest | 0.66 | 0.43 | 0.21 |
| UTE | No thrombin | Linear LASSO | -0.71 | -0.13 | 0.27 |
| | | Poly LASSO | -0.48 | -0.13 | 0.27 |
| | | Random Forest | -0.40 | -0.23 | 0.28 |
| | Thrombin | Linear LASSO | 0.14 | 0.02 | 0.27 |
| | | Poly LASSO | -0.24 | -0.03 | 0.28 |
| | | Random Forest | -0.20 | -0.16 | 0.30 |
| T1 FLASH | No thrombin | Linear LASSO | -0.25 | -0.10 | 0.30 |
| | | Poly LASSO | -0.42 | -0.43 | 0.34 |
| | | Random Forest | -0.18 | -0.28 | 0.32 |
| | Thrombin | Linear LASSO | 0.67 | 0.45 | 0.21 |
| | | Poly LASSO | 0.68 | 0.44 | 0.21 |
| | | Random Forest | 0.58 | 0.33 | 0.23 |
| Pooled Features | No thrombin | Linear LASSO | 0.79 | 0.61 | 0.18 |
| | | Poly LASSO | 0.63 | 0.39 | 0.22 |
| | | Random Forest | 0.89 | 0.75 | 0.14 |
| | Thrombin | Linear LASSO | 0.81 | 0.64 | 0.17 |
| | | Poly LASSO | 0.81 | 0.60 | 0.18 |
| | | Random Forest | 0.90 | 0.78 | 0.13 |

**Abbreviations:** RMSE = Root mean standard error; LASSO = least absolute shrinkage and selection operator

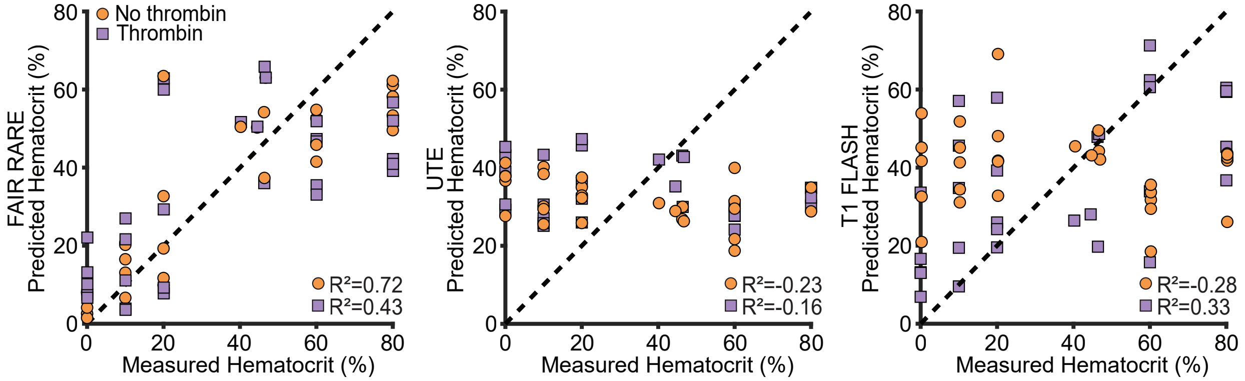


**Fig S3.** Random forest regression shows sequence-dependent variability in predicting clot hematocrit from radiomic features. Results using features from FAIR RARE, UTE, and T1 FLASH images are shown. Each data point represents a single clot. Data points are colored by condition: round orange indicates clots coagulated without thrombin, and square purple indicates clots coagulated with thrombin. The dotted line indicates perfect prediction (y = x). Regressions are based on 5-fold cross-validation.

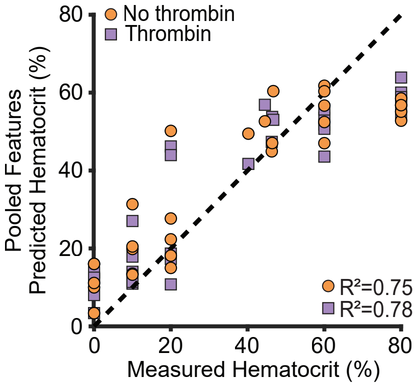


**Fig S4.** Random forest regression predicts clot hematocrit using pooled radiomic features from all nine MRI scans. Each data point represents a single clot. Data points are colored by condition: round orange indicates clots coagulated without thrombin, and square purple indicates clots coagulated with thrombin. The dotted line indicates perfect prediction (y = x). Regressions are based on 5-fold cross-validation.

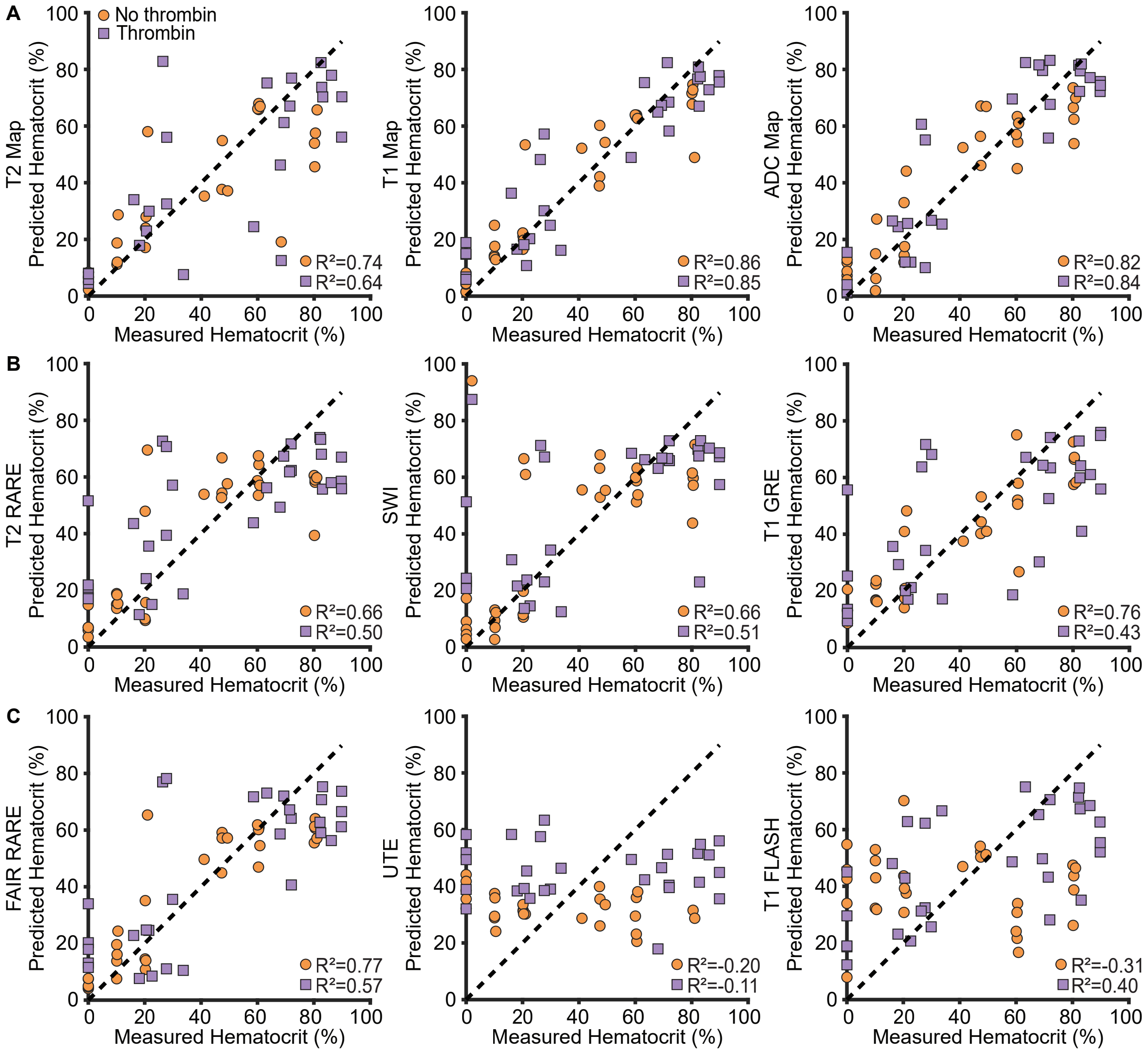


**Fig S5.** Random forest regression shows sequence-dependent variability in predicting corrected clot hematocrit from radiomic features. (A) Predictions based on radiomic features extracted from T2, T1, and ADC maps. (B) Predictions based on radiomic features from T2 RARE, SWI, and T1 GRE images. Each data point represents a single clot. Data points are colored by condition: round orange indicates clots coagulated without thrombin, and square purple indicates clots coagulated with thrombin. The dotted line indicates perfect prediction (y = x). Regressions are based on 5-fold cross-validation.

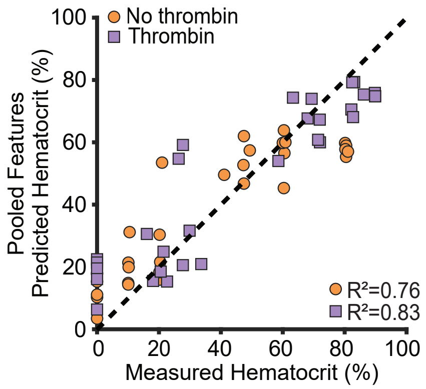


**Fig S6.** Random forest regression predicts corrected clot hematocrit using pooled radiomic features from all nine MRI scans. Each data point represents a single clot. Data points are colored by condition: round orange indicates clots coagulated without thrombin, and square purple indicates clots coagulated with thrombin. The dotted line indicates perfect prediction (y = x). Regressions are based on 5-fold cross-validation.

**Table S3.** Performance metrics for radiomic feature regression models predicting corrected hematocrit from individual MRI sequences and pooled features.

| Scan | Condition | Model | R | $R^2$ | RMSE |
|---|---|---|---|---|---|
| T2 Mapping | No thrombin | Linear LASSO | 0.80 | 0.58 | 0.18 |
| | | Poly LASSO | 0.67 | 0.45 | 0.21 |
| | | Random Forest | 0.86 | 0.74 | 0.15 |
| | Thrombin | Linear LASSO | 0.78 | 0.58 | 0.21 |
| | | Poly LASSO | 0.61 | 0.36 | 0.26 |
| | | Random Forest | 0.80 | 0.64 | 0.19 |
| T1 Mapping | No thrombin | Linear LASSO | 0.9 | 0.78 | 0.13 |
| | | Poly LASSO | 0.84 | 0.68 | 0.16 |
| | | Random Forest | 0.93 | 0.86 | 0.11 |
| | Thrombin | Linear LASSO | 0.88 | 0.76 | 0.16 |
| | | Poly LASSO | 0.88 | 0.75 | 0.16 |
| | | Random Forest | 0.93 | 0.85 | 0.12 |
| ADC Mapping | No thrombin | Linear LASSO | 0.59 | 0.33 | 0.23 |
| | | Poly LASSO | 0.38 | 0.14 | 0.26 |
| | | Random Forest | 0.91 | 0.82 | 0.12 |
| | Thrombin | Linear LASSO | 0.11 | 0.00 | 0.33 |
| | | Poly LASSO | 0.57 | 0.26 | 0.28 |
| | | Random Forest | 0.92 | 0.84 | 0.13 |
| T2 RARE | No thrombin | Linear LASSO | 0.72 | 0.50 | 0.20 |
| | | Poly LASSO | 0.71 | 0.50 | 0.20 |
| | | Random Forest | 0.82 | 0.66 | 0.17 |
| | Thrombin | Linear LASSO | 0.75 | 0.55 | 0.22 |
| | | Poly LASSO | 0.70 | 0.47 | 0.24 |
| | | Random Forest | 0.72 | 0.50 | 0.23 |
| SWI | No thrombin | Linear LASSO | 0.77 | 0.56 | 0.19 |

| | | | | | |
|---|---|---|---|---|---|
| | | Poly LASSO | 0.72 | 0.51 | 0.20 |
| | | Random Forest | 0.82 | 0.66 | 0.17 |
| | Thrombin | Linear LASSO | 0.76 | 0.58 | 0.21 |
| | | Poly LASSO | 0.77 | 0.58 | 0.21 |
| | | Random Forest | 0.71 | 0.51 | 0.23 |
| T1 GRE | No thrombin | Linear LASSO | 0.55 | 0.28 | 0.24 |
| | | Poly LASSO | 0.86 | 0.74 | 0.15 |
| | | Random Forest | 0.89 | 0.76 | 0.14 |
| | Thrombin | Linear LASSO | 0.72 | 0.49 | 0.23 |
| | | Poly LASSO | 0.70 | 0.47 | 0.24 |
| | | Random Forest | 0.66 | 0.43 | 0.25 |
| FAIR RARE | No thrombin | Linear LASSO | 0.81 | 0.63 | 0.17 |
| | | Poly LASSO | 0.81 | 0.63 | 0.17 |
| | | Random Forest | 0.88 | 0.77 | 0.14 |
| | Thrombin | Linear LASSO | 0.74 | 0.51 | 0.23 |
| | | Poly LASSO | 0.72 | 0.50 | 0.23 |
| | | Random Forest | 0.76 | 0.57 | 0.21 |
| UTE | No thrombin | Linear LASSO | -0.65 | -0.10 | 0.27 |
| | | Poly LASSO | -0.43 | -0.10 | 0.27 |
| | | Random Forest | -0.35 | -0.20 | 0.28 |
| | Thrombin | Linear LASSO | -0.52 | -0.06 | 0.33 |
| | | Poly LASSO | -0.27 | -0.04 | 0.33 |
| | | Random Forest | -0.04 | -0.11 | 0.34 |
| T1 FLASH | No thrombin | Linear LASSO | 0.01 | -0.27 | 0.32 |
| | | Poly LASSO | -0.42 | -0.41 | 0.34 |
| | | Random Forest | -0.10 | -0.31 | 0.33 |
| | Thrombin | Linear LASSO | 0.32 | -0.13 | 0.35 |
| | | Poly LASSO | 0.45 | 0.15 | 0.30 |
| | | Random Forest | 0.63 | 0.40 | 0.25 |
| Pooled Features | No thrombin | Linear LASSO | 0.80 | 0.63 | 0.17 |
| | | Poly LASSO | 0.67 | 0.44 | 0.21 |
| | | Random Forest | 0.89 | 0.76 | 0.14 |
| | Thrombin | Linear LASSO | 0.67 | 0.40 | 0.25 |
| | | Poly LASSO | 0.72 | 0.50 | 0.27 |
| | | Random Forest | 0.92 | 0.83 | 0.14 |

**Abbreviations:** RMSE = Root mean standard error; LASSO = least absolute shrinkage and selection operator

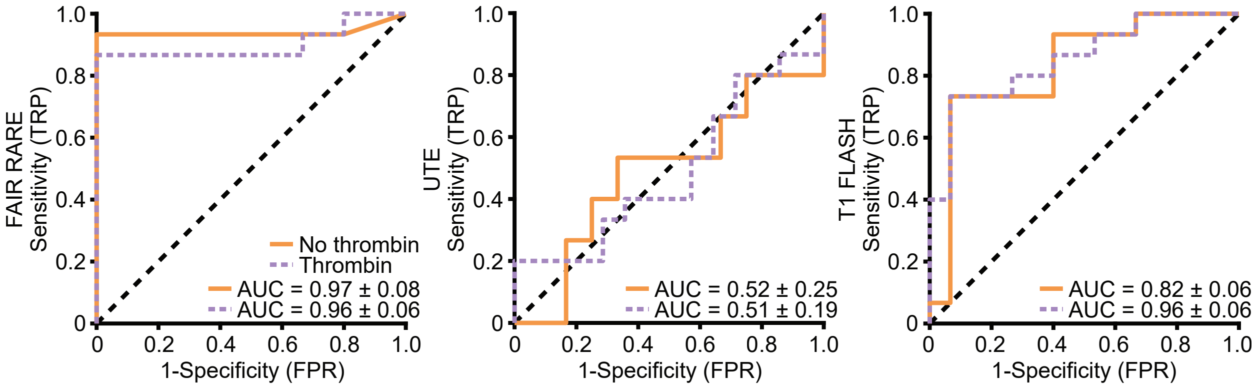


**Fig S7.** ROC analysis shows sequence-dependent performance in discriminating RBC-rich versus fibrin-rich clots using radiomic features from FAIR RARE, UTE, and T1 FLASH scans. Each curve represents the mean ROC across 5-fold cross-validation. Curves are colored by condition: orange indicates clots coagulated without thrombin, and purple indicates clots coagulated with thrombin. The area under the curve (AUC) $\pm$ standard deviation is shown on the plots.

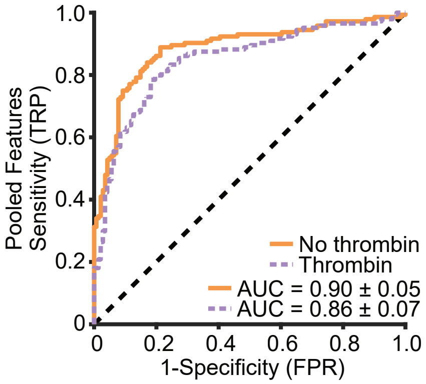


**Fig S8.** ROC analysis shows high performance in discriminating RBC-rich versus fibrin-rich clots using pooled radiomic features from all nine MRI scans. Each curve represents the mean ROC across 5-fold cross-validation. Curves are colored by condition: orange indicates clots coagulated without thrombin, and purple indicates clots coagulated with thrombin. The area under the curve (AUC) $\pm$ standard deviation is shown on the plot.

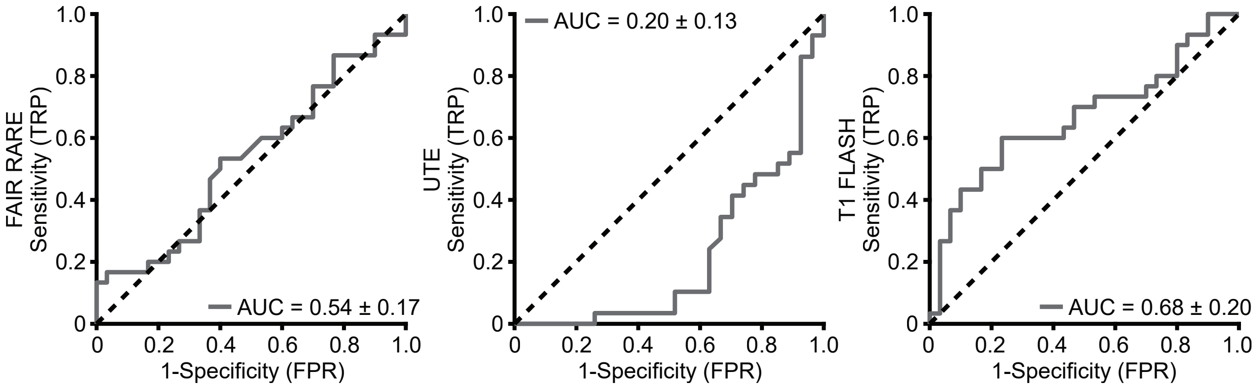


**Fig S9.** ROC analysis shows variable performance in discriminating clots coagulated with thrombin (contracted) versus without thrombin (uncontracted) using radiomic features from FAIR RARE, UTE, and T1 FLASH scans. Each curve represents the mean ROC across 5-fold cross-validation. The area under the curve (AUC) $\pm$ standard deviation is shown on the plots.

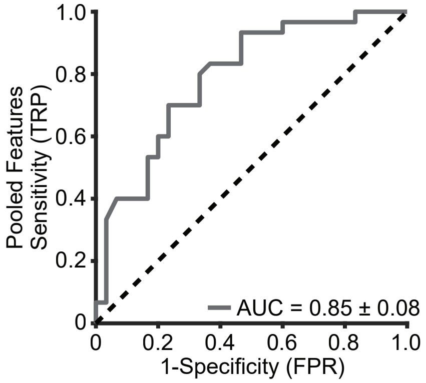


**Fig S10.** ROC analysis shows high performance in discriminating clots coagulated with thrombin (contracted) versus without thrombin (uncontracted) using pooled radiomic features from all nine MRI scans. The curve represents the mean ROC across 5-fold cross-validation. The area under the curve (AUC) $\pm$ standard deviation is shown on the plot.
